# An *in vivo* translation-reporter system for the study of protein synthesis in zebrafish embryos

**DOI:** 10.1101/336826

**Authors:** Inês Garcez Palha, Isabelle Anselme, Sylvie Schneider-Maunoury, François Giudicelli

## Abstract

Control of gene expression at the translation level is increasingly regarded as a key feature in many biological processes. Simple, inexpensive, and reliable procedures to visualise sites of protein production are required to allow observation of the spatiotemporal patterns of mRNA translation at subcellular resolution. We present a method, named SPoT (for Subcellular Patterns of Translation), developed upon the original TimeStamp technique (Lin et al., 2008), consisting in the expression of a fluorescent protein fused to a tagged, self-cleavable protease domain. Addition of a cell-permeable protease inhibitor instantly stabilizes newly produced, tagged protein allowing to distinguish recently synthesized protein from more ancient one. After a brief protease inhibitor treatment, the ratio of tagged *vs* non-tagged forms is highest at sites where proteins are the most recent, *i.e.* sites of synthesis. Therefore, by comparing tagged and non-tagged protein it is possible to spotlight sites of translation. By specifically expressing the SPoT cassette in neurons of transgenic zebrafish embryos, we reveal sites of neuronal protein synthesis in diverse cellular compartments during early development.

## INTRODUCTION

Organisms orchestrate their own gene expression program not only by the production of messenger RNA in appropriate amounts at the right time in the right cells, but also through the controlled translation of these mRNA molecules in time and space within the cells. Thus, subcellular patterns of protein synthesis are thought to play important roles in many biological processes such as early oocyte regionalisation, epithelial polarity and cell migration (reviewed in Medioni et al., 2012). Due to their large size and highly polarized shape, neurons are particularly sensitive to subcellular regulation of gene expression. Processes such as axon guidance, synapse formation or synaptic plasticity are dependent on external stimuli, which require swift responses in strictly defined cellular compartments (Holt and Schuman, 2013). Temporal and spatial constraints limit the immediate contribution of the cell body to these processes. Local translation within neurites is thus a manner to synthesize the required proteins in a limited time at a particular place, as a response to external signals. For local translation to occur, the transport of specific transcripts to these compartments is essential. mRNA localization information is usually located in its 3’UTR (Andreassi and Riccio, 2009). Several reports indicate that neurons are able to produce proteins within axons, even though elements of the protein synthesis machinery have been difficult to characterise in axonal compartments, presumably because of their singular morphology (Merianda et al., 2009; Willis et al., 2005). A recent study estimates that in neurons differentiated in vitro from ES cells, almost half of the neurite-enriched proteome is encoded by neurite-localized mRNA, suggesting that mRNA transport and local translation is a common mechanism for protein localization in neurites (Zappulo et al., 2017). However, direct data concerning the subcellular patterns of protein synthesis in whole living organisms are scarce. This is because few techniques exist that enable visualisation of protein synthesis in whole tissues. Classical metabolic labelling methods using labelled amino acid analogues are too promiscuous to allow cellular resolution, requiring sophisticated next generation developments such as Non Canonical Aminoacid Tagging (Hinz et al., 2013). In vivo imaging methods based on the bleaching and subsequent neosynthesis of fluorescent protein (Aakalu et al., 2001; Leung et al., 2006) are labor-intensive and prone to adverse effects of the bleaching procedure. Methods that detect solely nascent protein during the translation process (Tanenbaum et al., 2014) require heavy technology to reliably detect the faint signals, making their adaptation to whole organisms a most challenging task. We present here SPoT (Subcellular Patterns of Translation), a simple, inexpensive tool to visualise patterns of protein translation in neurons of the developing zebrafish embryo.

## MATERIALS & METHODS

### Cloning of the SPoT translation reporter constructs

The TimeStamp cassette comprising the NS3 domain of the hepatitis C virus (HCV) polyprotein, flanked by its NS4A/B specific cleavage sites and followed by an HA epitope coding sequence was obtained from the PSD95 plasmid (Lin et al., 2008). It was cloned in frame with Venus coding sequence in plasmid 5xUAS:myrVenus_3’UTR_chicken_ß-actin (Baraban et al., 2013). Chicken *ß-actin*-ΔZipcode and zebrafish *tubulin-β5* and *neuroD* 3’UTR sequences as described in (Baraban et al., 2013) were substituted to the original chicken *ß-actin* 3’UTR to obtain the corresponding plasmids. All the sequences were checked and are available on request.

### Fish handling and DNA injection

Zebrafish were raised and maintained as described (Kimmel et al., 1995). All our experiments were made in agreement with the european Directive 210/63/EU on the protection of animals used for scientific purposes, and the french application decree “Décret 2013-118”. The projects of our group have been approved by our local ethical committee “Comité d’éthique Charles Darwin”. The authorization number is 2015051912122771 v7 (APAFIS#957). The fish facility has been approved by the French “Service for animal protection and health”, with the approval number A-75-05-25. Embryos were staged according to the number of hours (hpf) or days (dpf) post-fertilization. Wild-type and huC:Gal4 (Akerboom et al., 2012) strains were used.

Translation reporter transgenic lines were obtained by injecting SPoT plasmid DNA in 1-2 cell stage huC:Gal4 zebrafish embryos with I-Sce meganuclease enzyme (New England Biolabs), in order to facilitate genomic integration. F0 embryos were first selected for GFP fluorescence at 1 dpf, then raised and selected for transgene transmission by crossing with wild-type mates.

### Danoprevir and cycloheximide treatment

Danoprevir (AdooQ Bioscience reference A10284) was resuspended in dimethylsulfoxyde (DMSO) and kept as a 10 mM stock solution at −20°C. Embryos were first dechorionated before treatment, then transferred to E3 medium containing freshly diluted Danoprevir at the desired concentration (40 µM for most experiments, unless specified otherwise). Danoprevir in egg water appeared not to present any toxicity for the zebrafish embryos: even at the highest concentrations used in our experiments (100 µM), embryos could be kept in the presence of the drug for up to several days and show no visible difference as compared to controls (not shown).

To inhibit protein synthesis, cycloheximide was diluted at 200 µg/ml in E3 medium in a similar manner.

All treatments were performed 28-30 hours after fertilisation (prim-15 stage).

### Whole-mount immunostaining

Embryos were fixed overnight at 4°C in 4% paraformaldehyde (PFA) in phosphate buffered saline (PBS), dehydrated in methanol at −20°C and then progressively rehydrated in PBS Tween (PBS with 0.1% Tween-20, Sigma). After rehydration, embryos were depigmented in 0.1M KOH/0.6% H2O2 in PBS until no pigmentation remained, then blocked for 1h at room temperature in blocking solution (0.1% BSA, 0.5% Triton X-100, 1% DMSO in PBS). They were incubated successively with primary antibodies recognizing Venus (GFP-1020 chicken polyclonal, Aves Lab ref, 1/500) and HA (3F10 mouse monoclonal, Sigma, 1/800), then with DAPI and secondary antibodies FITC-coupled anti-chicken IgY (Jackson ImmunoResearch 703-096-155, 1/800) and Alexa568-coupled anti-rat IgG (Molecular probes A-11077, 1/800), at least 24h each in blocking solution.

### Imaging

The yolk of immunostained embryos was manually removed and the embryos flat-mounted on microscope slides in 90% glycerol. Mounted embryos were imaged with a Nikon Eclipse E800 epifluorescence microscope, or optically sectioned on a Leica Sp5 confocal microscope.

## RESULTS AND DISCUSSION

### Design and validation of the SPoT method

In order to visualise sites of protein synthesis in the zebrafish embryo, we intended to develop a translation reporter system that would fulfil the two following requirements:

i. allow to easily distinguish newly synthesized proteins from more ancient ones
ii. be expressed specifically in defined cells through genetic control.

For this purpose, we combined the TimeStamp technique (Butko et al., 2012; Lin et al., 2008) with the reporter system for mRNA axonal transport previously developed in our lab (Baraban et al., 2013). The TimeStamp cassette encodes the proteolytic domain of the hepatitis C virus NS3 protease, flanked by its own specific cleavage sites and followed by a hemagglutinin (HA) epitope tag. Thus, the resulting fusion protein is tagged with a HA epitope that is quickly removed and degraded by default activity of the NS3 protease, unless a specific protease inhibitor is added, maintaining the tag fused to the protein of interest. This allows specific labelling of protein that has been synthesized after the addition of protease inhibitor, which is HA-tagged, as opposed to protein synthesised more anciently, which is not.

We transposed the TimeStamp cassette onto UAS:mVenus constructs, designed to express a myristoylated fluorescent protein (mVenus) in cells expressing the Gal4 transactivation factor (Baraban et al., 2013). We grafted the protease-HA tag cassette in frame at the 3’ end of the Venus coding sequence so that the product of translation would be both detected as fluorescent protein Venus and as HA-tagged, the latter representing only an unstable, short-lived form (Fig. 1A).

**Figure 1:**
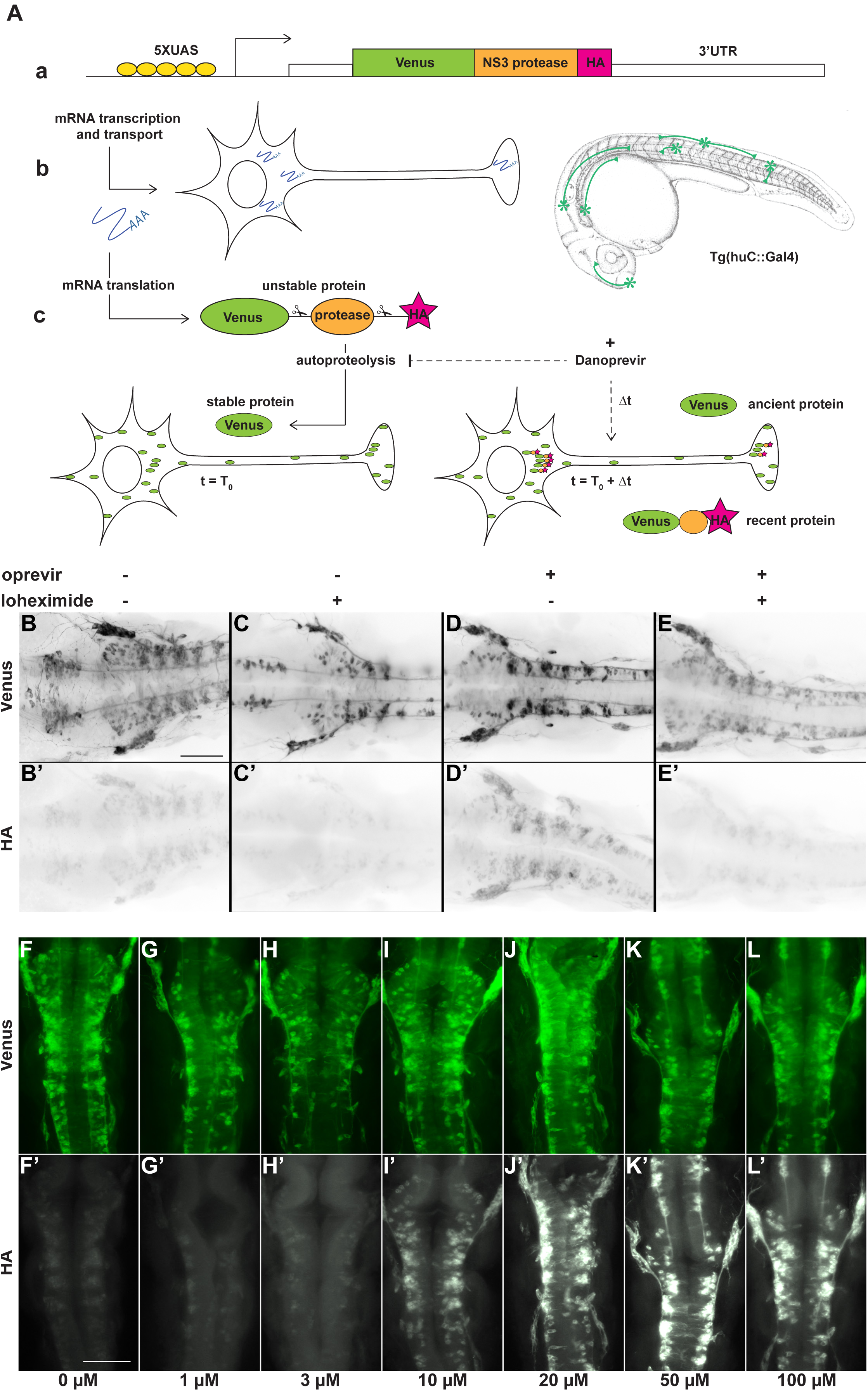
Protease-controlled translation reporter system. **(A)** Principle of the SPoT method. a) The translation reporter cassette encodes a myristoylated form of the Yellow Fluorescent Protein Venus fused to NS3 protease domain, flanked by two of its own specific cleavage sites, and a HA epitope tag. b) The UAS-controlled promoter ensures expression of the construct in neurons in Tg(huC:Gal4) transgenic line. c) In normal conditions, the protease cleaves itself rapidly after translation, removing the HA peptide. At time T0, addition of a small cell-permeable molecule (Danoprevir) that prevents protease activity stabilises subsequently synthesised proteins, maintaining their HA tag. HA-labelled protein, that have been synthesised during the Δt period between T0 and fixation, reveal sites of translation. **(B-E’)** Flatmounted immunostaining for Venus (top row) and HA (bottom row) of Tg(SPoT_3’ UTR chicken ß-actin) embryos incubated for 1h with Danoprevir (D,D’), Cycloheximide (C,C’), both (E,E’) or none (B,B’). **(F-L’)** Flatmounted immunostaining for Venus (top row) and HA (bottom row) of Tg(SPoT_3’ UTR chicken ß-actin) embryos incubated for 1h30 with increasing concentrations of Danoprevir. Scale bars: 200 µm.

To specifically express the construct in isolated neurons we used a huC:Gal4 zebrafish line, which expresses Gal4 in differentiated, immature neurons (Akerboom et al., 2012). Importantly, since 3’UTR sequences are known to influence mRNA fate, we generated SPoT constructs and transgenic lines with various 3’UTRs, initially concentrating on 3’UTRs that we had previously showed to allow axonal transport, chicken *ß-actin* and zebrafish *tubulin-β5* (Baraban et al., 2013).

When injected in heterozygous huC:Gal4 eggs at 1-2 cell stage, the UAS:SPoT constructs yielded robust Venus fluorescence in isolated neurons, with highly variable level of expression among embryos, a common feature in transient transgenesis approaches. In order to get more reproducible and even expression levels, we created transgenic lines of zebrafish having integrated the construct in their genomes, and studied its expression in their progeny. As expected, the stable transgenic embryos exhibited more uniform levels of expression in primary neurons (Fig. 1B), although sometimes subject to variegation (silencing in a subset of the targeted cells), a feature quite common in zebrafish transgenic embryos.

### The SPoT product is properly self-cleaved in zebrafish embryos

We first sought to assess the efficiency of autoproteolysis of the SPoT protein in zebrafish neurons, by attempting to detect HA immunoreactivity in untreated SPoT transgenic embryos by immunofluorescence. As shown in Fig. 1B’, we consistently detected a weak, but specific HA staining in Venus-expressing neurons. This contrasts with the original Timestamp publications, where the tag epitope was self-cleaved rapidly enough so that it was not detectable in control conditions (Lin et al., 2008). To test whether this weak HA signal represented transient, recently produced protein that had not yet been cleaved, or if it resulted from imperfect self-cleavage of the SPoT protein, we treated embryos with the protein synthesis inhibitor cycloheximide for various durations before fixation and immunostaining. We observed that inhibition of protein synthesis progressively reduced the HA signal, and that 1 hour treatment was sufficient to essentially suppress this residual HA signal in the embryos (Fig. 1C,C’). This demonstrates that the residual HA staining in control embryos consisted solely of protein that had been synthesized within the last hour before fixation, rather than a subpopulation of SPoT protein that would be resistant to self-cleavage.

This residual signal in untreated controls was not described in the original TimeStamp publication (Lin et al., 2008). This could be due to differences between the experimental systems, such as more intense translation activity in zebrafish embryonic neurons than in mouse cultured neurons, or to intrinsically lessened activity of the NS3 protease in zebrafish cells. The former explanation is supported by the fact that, in zebrafish SPoT embryos compared to TimeStamp cultured cells, shorter exposures to the protease inhibitor are sufficient in order to obtain robust signal (45 minutes vs 2 hours). In addition, although one could have expected the activity of a protease, which normally operates at 37°C, to depend on temperature and possibly decline at 28°C, where zebrafish eggs normally develop, it appears not to be the case: we observed no difference in residual HA signal in untreated embryos grown at low (23 °C), normal (28°C), or high (33°C) temperatures (data not shown).

### Protease inhibitor treatment highlights newly synthesized proteins

Even though HA staining in control embryos designated a fraction of recently synthesised protein that had not yet been self-cleaved, this fraction was clearly too small to be of use for our reporter purpose, as stronger signal was required. We then tested whether inhibition of protease activity could reveal sites of translation by stabilising newly synthesised HA-tagged SPoT protein.

We used Danoprevir, a known inhibitor of NS3 protease (Jiang et al., 2014), to inhibit self-cleavage of the SPoT protein during a controlled time span preceding fixation, thereby highlighting protein synthesized during this period.

Upon 1h incubation with 40 µM Danoprevir, Tg(SPoT) embryos exhibited marked accumulation of HA immunoreactivity throughout Venus-positive neurons (Fig. 1D,D’), showing that this small molecule effectively blocks autoproteolysis of the SPoT protein product.

To confirm that the accumulated HA signal upon Danoprevir treatment truly represented *de novo* protein synthesis, we inhibited protein synthesis during Danoprevir treatment by adding cycloheximide simultaneously. In that case, we observed a result similar to treatment with cycloheximide alone, with barely detectable HA signal (Fig. 1E,E’). This confirms that accumulation of HA signal upon Danoprevir treatment is entirely imputable to *de novo* synthesis of the TimeSTAMP protein during the time of treatment.

To determine which dose of protease inhibitor resulted in optimal labelling of newly synthesized SPoT protein, we treated Tg(SPoT) embryos for 1h30 with increasing concentrations of Danoprevir. While HA signal remained faint at Danoprevir concentrations of 1 or 3 µM (Fig. 1F-H’), it markedly accumulated at 10 and 20 µM (Fig. 1I-J’), and reached a plateau at 50 and 100 µM (Fig. 1K-L’). Therefore, concentrations between 40 and 80 µM seem to saturate the cleavage sites and maximize SPoT protein stabilisation. These saturating concentrations were used in all subsequent experiments.

The cell-type specificity obtained by the Gal4-UAS transactivation system ensures that the observed accumulation actually represents protein synthesis in the neurons themselves and cannot be due, as has been suggested in other contexts, to intercellular transfer from neighbouring glia. Furthermore, it may permit the use of the method in any cell type for which there exists specific Gal4 transgenic lines, although the validity in other contexts would have to be confirmed by performing appropriate controls such as cycloheximide inhibition of protein synthesis.

### Visualising subcellular sites of translation

We then proceeded to observe whether accumulation of newly synthesised TimeStamp protein allowed to characterise particular sites of translation. To this end, we processed Danoprevir-treated embryos for HA and Venus whole-mount immunostaining, then optically sectioned the embryos and systematically compared HA signal (recent protein) to Venus signal (total protein). In these images, Venus staining typically extended over the whole span of individual neurons, including distant extensions (Fig. 2A). In contrast, HA staining, representing the most recently synthesised fraction of the SPoT protein, appeared concentrated in defined areas, thereby identified as sites of translation (Fig. 2A’).

**Figure 2:**
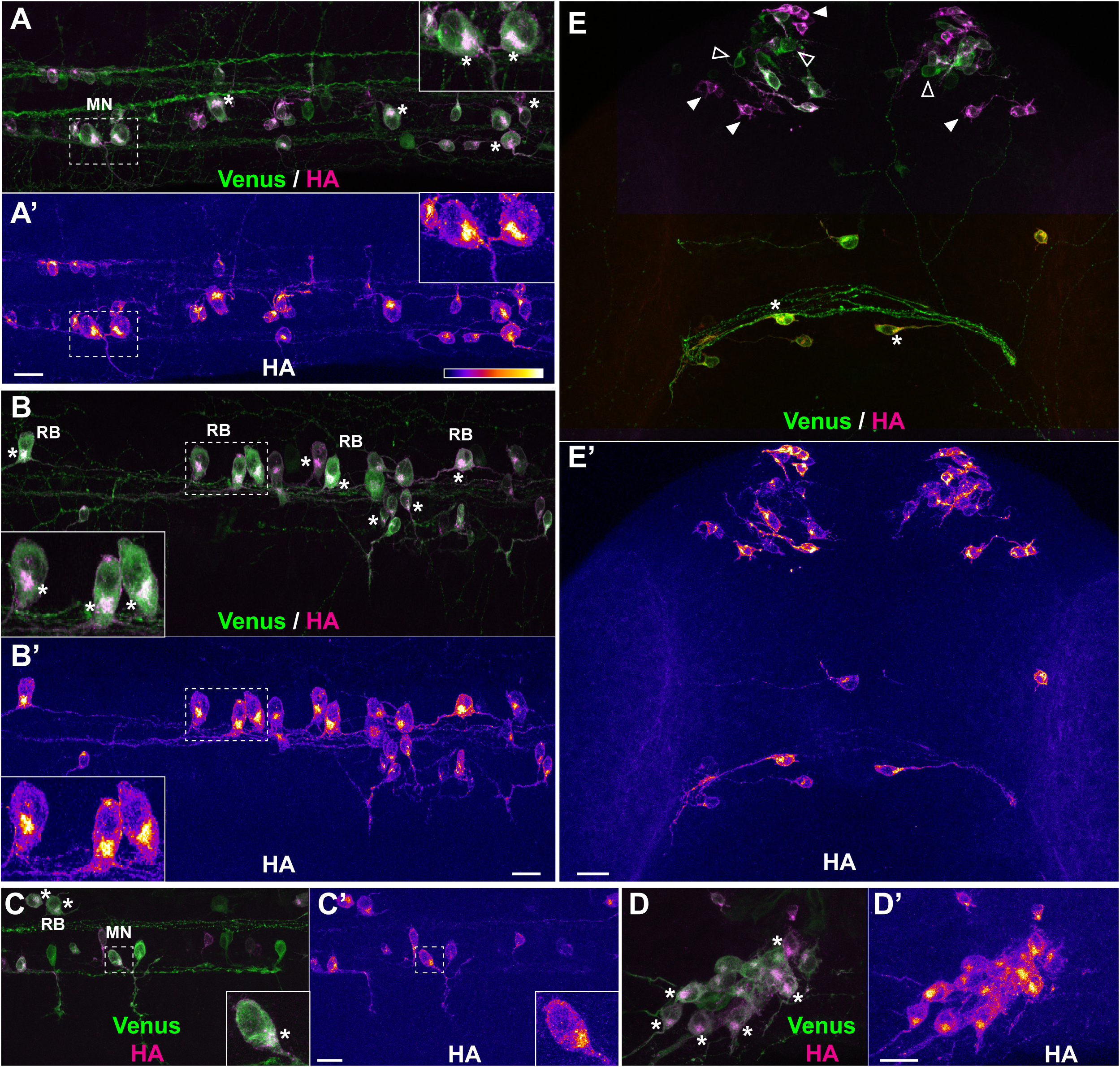
Short treatment with protease inhibitor highlights sites of protein synthesis. Representative examples of protein synthesis patterns revealed by HA/Venus immunostaining of Danoprevir-treated embryos. (**A-E)** Superimposed Venus (green) and HA (magenta) immunostaining signals. **(A’-E’)** HA immunostaining signal intensity color-coded with imageJ ‘Fire’ lookup table. Colour-intensity correspondence is represented in the calibration bar. **(A-C)** Lateral views of the trunk region. **(D)** Dorsal view of the trigeminal ganglion. **(E)** Dorsal view of the anterior head region. **(A,B,E)** Transgenic line = Tg(SPoT_3’ UTR tubb5). Danoprevir treatment time = 2h **(C,D)** Transgenic line = Tg(SPoT_3’ UTR chicken ß-actin). Danoprevir treatment time = 1h30 Scale bars: 20 µm. Insets represent 2-fold (A,B) or 3-fold (C) magnifications of the dashed boxes. Asterisks highlight cells with marked accumulation of newly synthesised protein between the nucleus and axon initial segment. In (E), plain white arrowheads indicate neurons with high newly synthesised SPoT protein, while empty arrowheads indicate neurons which express the SPoT transgene but have not produced SPoT protein during the time of treatment. MN: Motor neuron. RB: Rohon-Beard sensory neuron. pc: posterior commissure.

In zebrafish primary neurons, the most conspicuous sites of translation revealed by this technique was the portion of the cell body situated between the nucleus and the axon initial segment (asterisks in Fig. 2A-E). This was particularly evident in large neurons, such as spinal cord sensory (Rohon-Beard, marked RB in Fig. 2B,C) or motor (marked MN in Fig. 2A,C) neurons, or peripheral sensory neurons of the trigeminal ganglion (Fig. 2D). We also observed that among neurons seemingly expressing similar levels of SPoT protein (as assessed by Venus staining) some could have been actively producing the protein during the time of Danoprevir treatment (high HA staining, plain arrowheads in Fig. 2E), while their neighbours had produced very little, if any, during the same period (empty arrowheads in Fig. 2E; see also Fig. 3B and 3G). This indicates that protein synthesis likely occurs in phases in these neurons, with bursts of expression alternating with troughs in protein synthesis activity. Whether these phases correspond to cell-wide metabolic phases of protein synthesis or whether they reflect transcript-specific variations in translation efficiency with time could not be determined with the SPoT technique.

**Figure 3:**
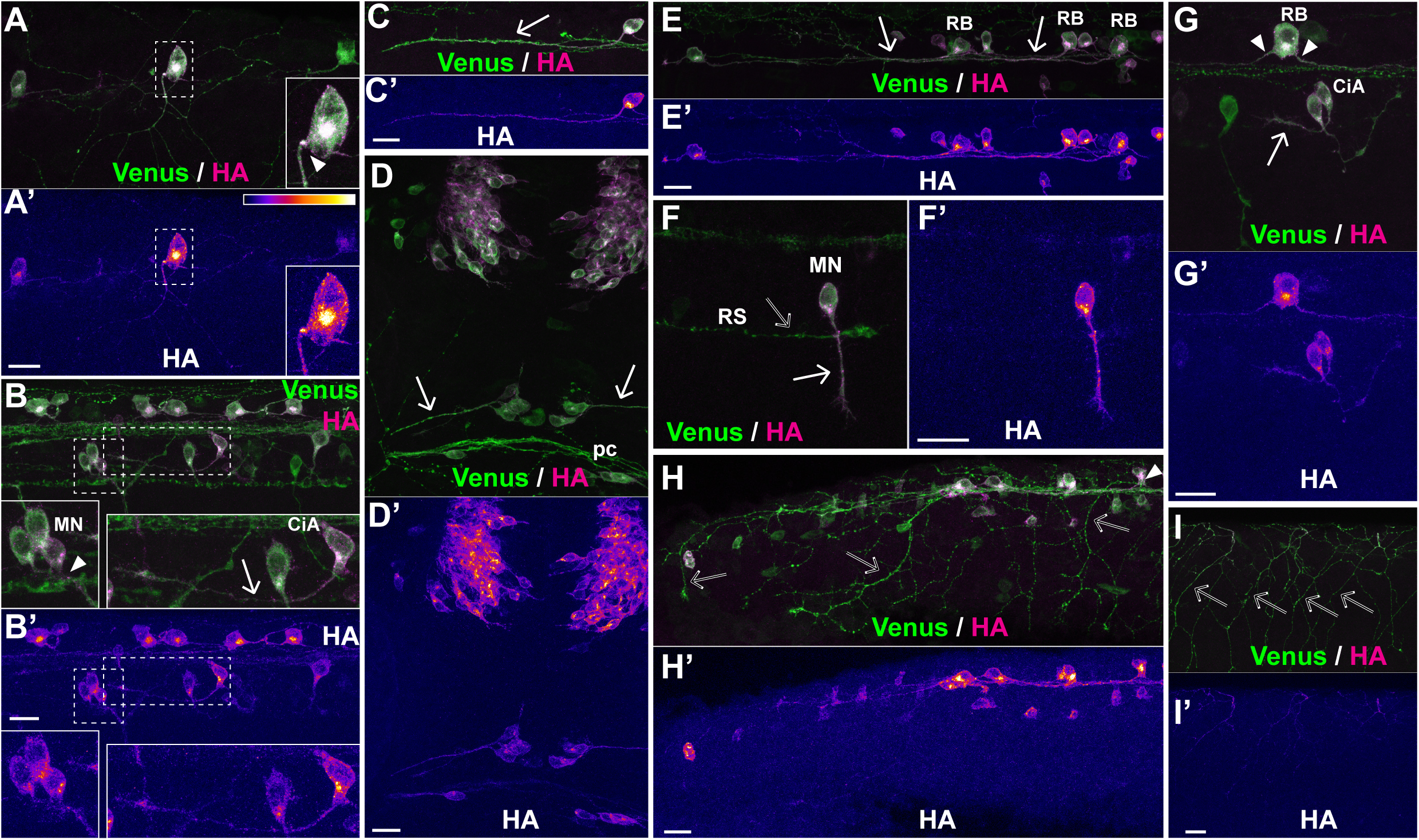
The SPoT reporter reveals translation sites along axons. Representative examples of sites of protein synthesis within axons revealed by HA/Venus immunostaining of Danoprevir-treated embryos. **(A-I)** Superimposed Venus (green) and HA (magenta) immunostaining signals. **(A’-I’)** HA immunostaining signal intensity color-coded with imageJ ‘Fire’ lookup table. Colour-intensity correspondence is represented in the calibration bar. **(A,B,E)** Lateral views of the trunk region. **(C,H)** Lateral views of the tail region. **(D)** Dorsal view of the anterior head region. **(F)** Isolated spinal motor neuron; the distal part and growth cone of a large reticulospinal axon is visible, but has no HA staining. **(G)** Lateral view of isolated RB, motor and inter neurons of the spinal cord. **(I)** Peripheral arbours of sensory neurons innervating the skin. Note the complete absence of HA signal. Arrowheads mark sites with characteristic accumulation of newly synthesised SPoT reporter in axon initial segment. Arrows point to axons with newly synthesised protein all along the shaft length. Empty arrows point to prominent axons with no protein synthesis. CiA: Circumferential Ascending neuron. MN: Motor neuron. RB: Rohon-Beard neuron. RS: Distal portion of a reticulospinal projection. **(A,B,D,F,H,I)** Transgenic line = Tg(SPoT_3’ UTR chicken ß-actin). Danoprevir treatment time = 1h30 **(C,E)** Transgenic line = Tg(SPoT_3’ UTR tubb5). Danoprevir treatment time = 2h **(G)** Transgenic line = Tg(SPoT_3’ UTR tubb5). Danoprevir treatment time = 40 minutes. Scale bars: 20 µm. Insets represent 2-fold magnification of the dashed boxes.

### Protein synthesis in axons

Since protein synthesis in axons, formerly presumed not to occur (see Alvarez et al., 2000 for a review), has now attracted considerable attention as a possible mechanism underlying axonal growth, survival, and guidance (recent reviews: Cioni et al., 2018; Riccio, 2018), we particularly scrutinised whether the SPoT transgene allowed detecting translation events in axons. In the SPoT transgenes used above, the SPoT protein is produced from a mRNA containing the 3’UTR sequence of chicken beta-actin (SPoT_3’UTR ß-actin) or tubulin-β5 (SPoT_3’UTR tubb5) mRNA, both of which allow axonal transport in zebrafish neurons (Baraban et al., 2013). Therefore, we searched whether Danoprevir treatment enabled detecting axonal translation of the reporter. We observed considerably less accumulation of newly synthesised HA-tagged protein in axons compared to the aforementioned accumulations in neuronal somata, indicating, perhaps not surprisingly, that the bulk of translation events of the SPoT mRNA occurs in the cell body. Nevertheless, we found that some, although not all, axons did exhibit focal accumulation of newly synthesised HA-tagged proteins upon incubating transgenic embryos with Danoprevir (Fig. 3). In agreement with previously observed localisation of mRNA (Baraban et al., 2013), newly synthesised protein frequently accumulated in the axon initial segment (arrowheads in Fig. 3A,B,G,H). But we also observed newly synthesised protein in axon shafts (plain arrows in Fig. 3C,D,F) and at the growth cone (Fig. 3F). Notably, this accumulation was specific to some cell types. Typically, spinal motor neurons (Fig. 3F) or circumferential ascending (CiA; Bernhardt et al., 1990) interneurons (Fig. 3B,G) often exhibited axonal protein synthesis. In contrast, we never observed foci of protein synthesis in peripheral sensory projections (empty arrows in Fig. 3H-I’) or reticulospinal projections in the medial longitudinal fasciculus (mlf) (empty arrow in Fig. 3F,F’).

In order to determine whether the observed axonal translation of the reporter construct correlated with transport of its mRNA to axons, we generated two additional zebrafish transgenic lines where the 3’UTR of the SPoT cassette had been substituted either with the 3’UTR sequence of zebrafish *neuroD* gene, which encodes a neuronal transcription factor, or with a version of the *ß-actin* 3’UTR deleted in the axonal localisation zipcode (Kislauskis et al., 1994). We showed previously that when included in reporter constructs, these 3’UTR restrict axonal transport (Baraban et al., 2013).

When SPoT_neuroD (Fig. 4A-B’) or SPoT_chicken_ß-actin-ΔZipcode (Fig. 4C,C’) transgenic embryos were treated with Danoprevir for 1h30, we observed accumulation of the HA signal in neurons, denoting active translation of the reporter during the time of treatment. Interestingly however, there was much less accumulation in distal axons than in SPoT_chicken_ß-actin transgenics, presumably reflecting limited axonal translation consecutive to restricted axonal transport of mRNA.

**Figure 4:**
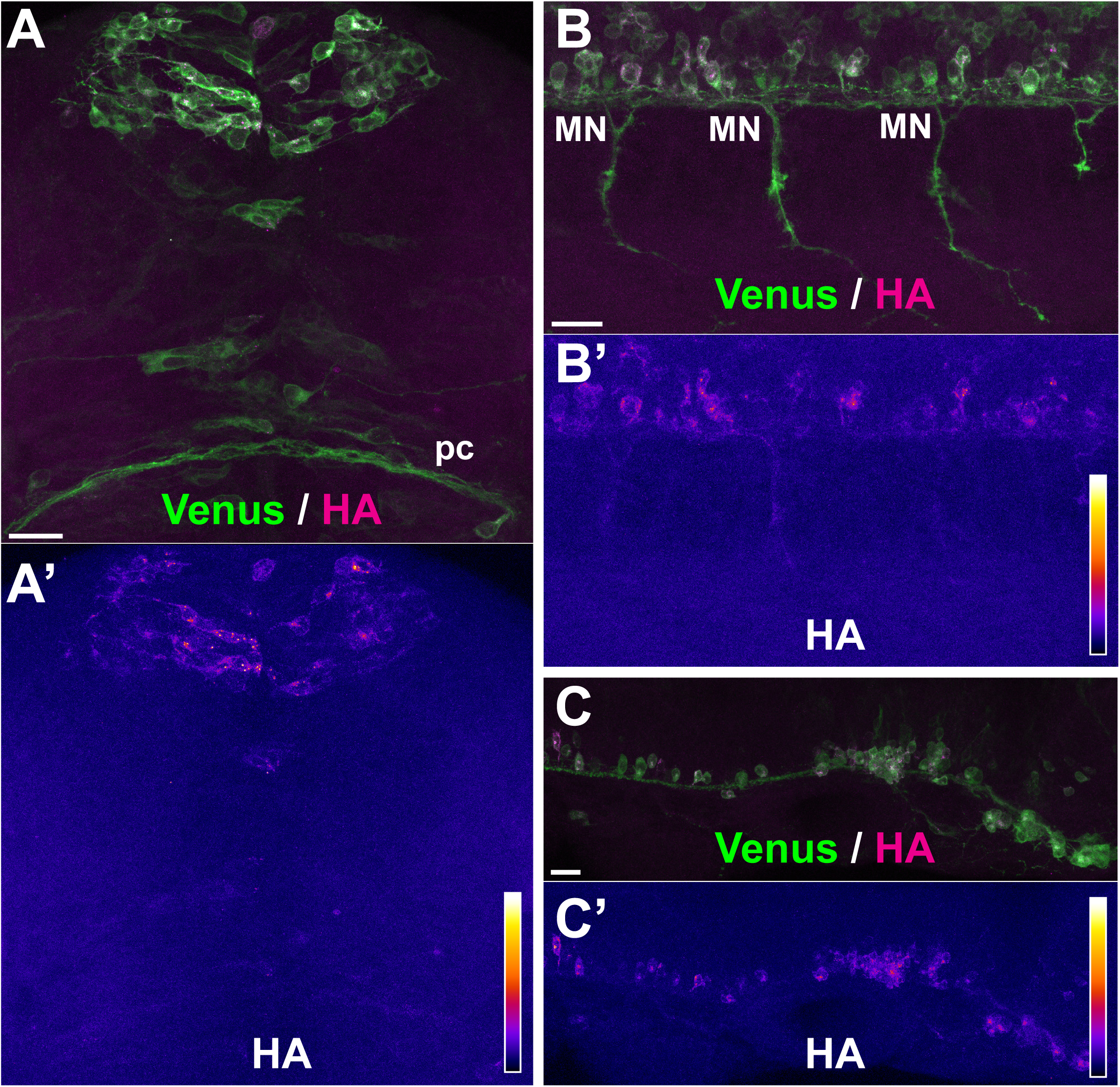
No axonal translation occurs with soma-restricted 3’UTR sequences. Representative examples of sites of protein synthesis revealed by HA/Venus immunostaining of Danoprevir-treated, SPoT transgenic embryos containing 3’UTRs with restricted axonal mRNA localisation. **(A-C)** Superimposed Venus (green) and HA (magenta) immunostaining signals. **(A’-C’)** HA immunostaining signal intensity color-coded with imageJ ‘Fire’ lookup table. Colour-intensity correspondence is represented in the calibration bar. **(A,B)** Transgenic line = Tg(SPoT_3’ UTR neuroD). Danoprevir treatment time = 1h30 **(C)** Transgenic line = Tg(SPoT_3’ UTR chicken ß-actin-ΔZipcode). Danoprevir treatment time = 1h30 Scale bars: 20 µm

In conclusion, the SPoT translation reporter system is a simple tool for visualising translation patterns in neurons and possibly other cell types within whole embryos. Here, it allows us to identify sites of protein synthesis in axons and growth cones, in several neuronal types while they are absent from others. By design, SPoT requires specimen to be fixed prior to analysis, therefore cannot follow protein synthesis in real time, unlike other, more sophisticated, techniques (SunTag, FUNCAT). Nevertheless, it provides a comprehensive view of protein synthesis events across whole embryos. Therefore, the SPoT method may allow recurrent questions about where and when protein is synthesised within cells to be explored more systematically.

## ACKNOWLEDGEMENTS

The authors wish to thank Alex Bois, Stéphane Tronche, and all staff of the IBPS aquatic facility for outstanding fish care.

This work was supported by the Centre National de la Recherche Scientifique (CNRS), the Institut National de la Santé et de la Recherche Médicale (Inserm), the Université Pierre et Marie Curie (now Sorbonne Université), and grants from the Agence Nationale pour la Recherche (ANR “RNAGRIMP”, FG partner) and Fondation pour la Recherche Médicale (FRM DEQ20140329544 to SSM). IP was supported by a PhD contract from Sorbonne Université.

